# Method for Modeling Residual Variance in Biomedical Signals Applied to Transcranial Doppler Ultrasonography Waveforms

**DOI:** 10.1101/633669

**Authors:** Kian Jalaleddini, Samuel G. Thorpe, Nicolas Canac, Amber Y. Dorn, Corey M. Thibeault, Seth J. Wilk, Robert B. Hamilton

## Abstract

*Transcranial Doppler* (TCD) ultrasonography measures pulsatile cerebral blood flow velocity in the arteries and veins of the head and neck. The velocity pulse waveform morphology has been shown to have physiological and diagnostic significance. However, the measured pulses may exhibit a high degree of variability that deteriorates the estimates of clinical parameters. This study characterizes the TCD residual variance that result in pulse variability.

We retrospectively utilized the data from 82 subjects. A trained sonographer insonated the middle cerebral arteries using a 2MHz hand-held probe. We implemented a multi-stage algorithm to identify the TCD residuals in each scan: pulses were identified; outlier pulses were flagged and removed; the average pulse waveform was taken as the ensemble average of the accepted pulse waveforms; finally, the resampled average pulse waveforms subtracted from individual pulses were taken as the TCD residuals. For each scan, we reported the signal to noise ratio and parameterized models for residuals: their amplitude structure using probability density function models and their temporal structure using autoregressive models.

The signal to noise ratio 90% range was [1.7, 18.2] dB. The estimated probability density functions were best characterized by a generalized normal distribution whose beta parameter was smaller than 2 in 93% of scans. The identified frequency structure showed the dynamics were low-pass in nature.

Analysis of the TCD residuals is useful in the assessment of the signal quality. Moreover, our identified models can also be used to generate synthetic TCD signal that enables future realistic simulation studies.

## 1. Introduction

*Transcranial Doppler* (TCD) is a diagnostic ultrasound-based imaging modality for real-time monitoring of the brain vasculature function through measurement of the *Cerebral Blood Flow Velocity* (CBFV) [1, 2]. CBFV is a pulsatile flow, thus, a number of features extracted from CBFV pulse morphology have been shown to have diagnostic significance such as *Pulsatility Index* (PI), *Thrombolysis In Brain Ischemia* (TIBI), *P2 Ratio* (P2R), and *Velocity Curvature Index* (VCI) [3, 4, 5, 6].

TCD measurement of the pulse waveform can exhibit variability that degrades the confidence of the estimated features and possibly the diagnosis. It is important to characterize the variability as a measure of signal quality and to understand how the TCD features are affected by the presence of variability in the CBFV waveform.

The variability within the measured TCD signal can have a number of origins. Intrinsic interpulse variability within measured TCD pulses can be due to healthy or pathologic arrhythmic heart rate variability, or pulse-level changes in the waveform morphology, e.g. changes in systole, diastole, P2R, etc [7, 8]. Another source of variation is changing the impedance between the TCD probe and the skull due to the movement of the patient or sonographer. Ambient noise, as well as signal processing artifacts generated in processing of the echoes, may also contribute to the observed variability [9].

The purpose of this study is to characterize the TCD residuals defined as the unmodeled dynamics in a TCD recording calculated from inter-pulse variability. This would be instrumental in the assessment of the measured signal quality and identification of confidence in estimated diagnostically-used TCD parameters. Moreover, such characterization can help generate realistic synthetic TCD signals that enable further simulation studies. In this study, we use a rich dataset comprising data from healthy, acute stroke patients, and in-hospital control subjects. We present an algorithm to create statistical models from TCD residuals and an algorithm to create random realizations from these models.

## 2. Methods

In this section, we describe our experiment protocol, discuss the algorithm to extract TCD residuals, and finally present the type of models we used to parameterize the residuals. Note that in the following, residuals refers to TCD residuals unless otherwise stated.

### 2.1. Experiments

For this study, we retrospectively analyzed the TCD data from subjects recruited at the Erlanger Health Systems Southeast Regional Stroke Center in Chattanooga, TN.

#### 2.1.1. Subjects

A trained sonographer acquired TCD waveforms from 82 subjects. The subjects were either diagnosed with *Large Vessel Occlusion* (LVO) of intracranial arteries confirmed with *Computed Tomography Angiography* (CTA) or in-hospital control subjects who arrived at the hospital presenting with stroke symptoms, but were later confirmed negative for LVO by CTA imaging. They also performed a follow-up TCD scan on the LVO subjects within 72 hours of injury. The experiment protocols were approved by the University of Tennessee College of Medicine Institutional Review Board (IRB# 16-097).

#### 2.1.2. TCD Scans

At each session, a trained sonographer acquired multiple TCD scans using a 2 MHz hand-held probe. They identified CBFV signals associated with the left or right *Middle Cerebral Arteries* (MCAs) by insonating through the transtemporal windows and obtained scans for as many depths as possible between 45-60 mm. Once an optimized CBFV signal with a smooth fitting envelope trace was identified, a 30 s scan began. Before analysis, we rejected the scans in which the CBFV was zero more than 10% of the scan time. In total, we included 2489 scans for this study. The sampling frequency was 125 Hz.

### 2.2. TCD Residuals Identification

We designed and implemented a multi-stage algorithm to identify residuals and created models for them. The algorithm first identifies an empirical model for the CBFV pulse waveform and takes residual as any deviation from this model. Thus, it is based on the assumption that the vascular activity generating the CBFV is time-invariant and so CBFV is periodic. Below, we describe these steps in detail:

**1- Pulse Detection:** First, the *Moving Difference Filter* (MDF) pulse detection algorithm identified the pulse start and stop times. This algorithm is described in detail in [10] and its accuracy is validated on a rich dataset consisting of healthy and pathologic CBFV waveforms. In summary, it consists of the following primary steps: (*i*) the CBFV signal is band-pass filtered to remove low-frequency and high-frequency noise; (*ii*) a moving difference filter is applied to enhance the upslope that defines the start of a typical CBFV pulse; (*iii*) a search window is defined on the CBFV signal to identify candidate pulse start times; (*iv*) pulse length and pulse alignment analyses are used to handle possible errors by fine-tuning the precise locations of the pulse start times.

**2- Pulse Rejection:** Second, the pulse rejection algorithm identified the pulses that were statistically different in a scan and removed them. This algorithm is described in detail in [11] and validated on a rich dataset consisting of healthy and pathologic CBFV waveforms. In summary, this algorithm works by comparing the correlation between individual pulses and the average pulse waveform and flagging those that statistically appear as outliers. To avoid possible biases in calculation of the statistics, the algorithm is iterative and removes the most prominent outlier, if any, at each iteration and recalculates the average pulse waveform from the accepted pulses up to that iteration.

**3- *Average Pulse Waveform* (APW):** Third, the APW was calculated. To this end, the average pulse length is calculated as the average length of the accepted pulses. Next, all the accepted pulses are normalized to the average pulse length. This is achieved by padding the short pulses with their last values and truncating the end of long pulses. The APW is then the ensemble average of the length-normalized pulses.

**4- TCD Residuals:** Finally, for every accepted pulse, the length of the APW was scaled to match the length of the pulse. This was achieved by using the resampling tool implemented in Scipy Signal Processing that transforms the signal to the frequency domain and then zero-pads or truncates the transformed signal to match the resampled length and then transform the signal back to the time domain [12]. The resampled APW was subtracted from the pulse to give the residuals. See Figure 1 for illustration of three examples.

**Figure 1:**
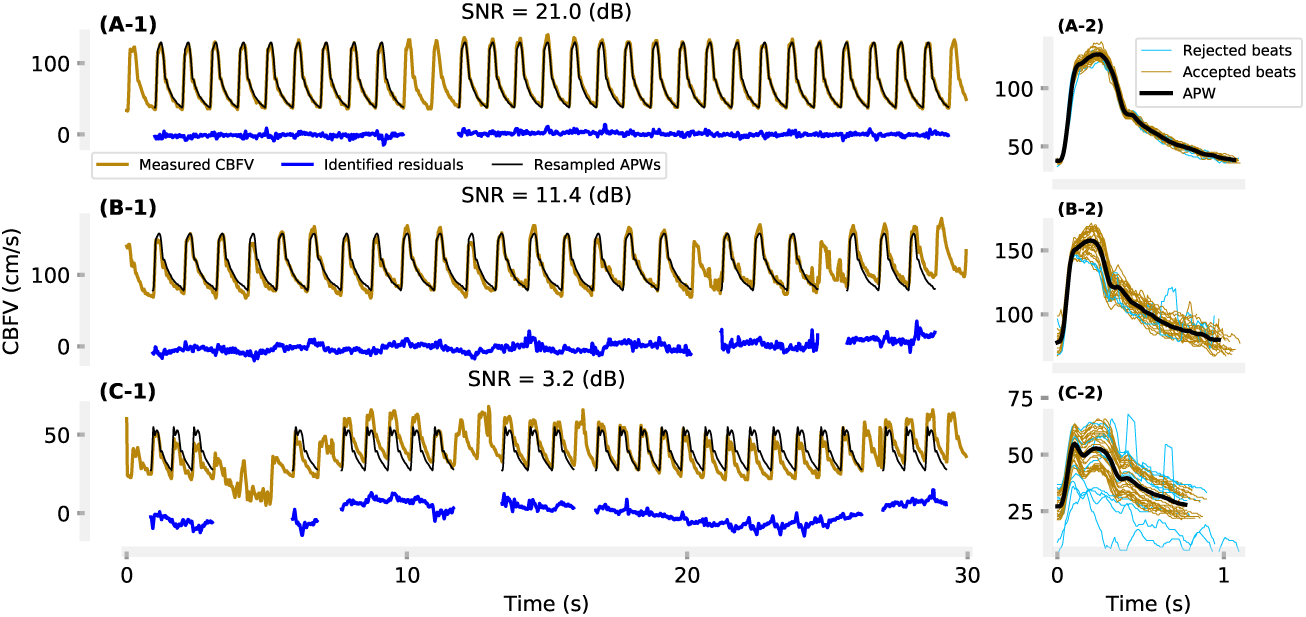
Three representative TCD scans with high, medium, and low SNR levels: (A-1, B-1, C1) measured TCD along with the predicted residuals and tiled replicated average pulse waveform; (A-2, B-2, C-2) ensemble of accepted pulses together with the average pulse waveform.

#### 2.2.1. Models of Residuals

Prior to the following modeling attempts, we removed the mean of the residuals.

##### Power

We expressed the power of the residuals in terms of *Signal to Noise Ratio* (SNR):

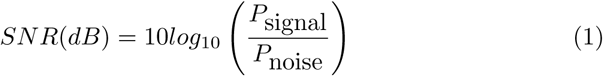

where *P* is power defined as the sum of the squared samples of a signal. “Noise” is the residuals and “signal” is the tiled replicated re-sampled APWs. We subtracted the means of the “signal” and “noise” for this calculation.

Since SNR defines the relative power of the TCD residuals, it can be served as an objective measure of signal quality and a meaningful metric characterizing the inter-pulse variability in a TCD recording.

##### Amplitude Structure

We calculated empirical *Probability Density Functions* (PDFs) with 20 bins for the residuals extracted from each TCD scan. To parameterize the distribution, we took a brute force approach. We fit more than 95 well-known continuous distributions, all those implemented in SciPy Statistical Functions version (scipy.stats) 1.1.0, to the residuals [12]. We used the SciPy fit function that returns the maximum likelihood function for parameters (shape, location, scale, etc.) of a distribution from the computed empirical distributions. This optimization does not necessarily have a closed form and typically the local optimal parameters are identified using numerical optimization algorithms. We chose the distribution with the highest average correlation between the empirical and theoretical PDFs across all the scans as the most likely model for the amplitude structure of the residuals.

##### Temporal Structure

The identified residuals can be modeled as a random process. Thus, we fit an Auto-Regressive (AR) model to the identified residuals of each scan to model their temporal structure. We chose the AR model because it is simple and easy to identify as the estimation of the parameters can be achieved using ordinary least-squares and this model can be easily simulated to create synthetic data.

### 2.3. Synthetic TCD Residuals Generation

In many studies of neurosonology, it is important to simulate TCD signals to test algorithms or train sonographers [13, 14, 15]. The above-identified models provide a means of generating synthetic TCD residuals. Thus, the goal is to generate realizations of non-white, non-Gaussian random sequences conforming to the identified models for distribution and spectral density. To this end, one can follow the following steps: (*i*) generate a sample for the parameter *β* from the distribution characterized in Figure 5; (*ii*) select an AR model from the ensemble of the identified AR models; (*iii*) generate a realization of stochastic time series from generalized normal distribution with the chosen scale parameter *β* and temporal characteristics according to the selected AR model (see details below); (*iv*) select and SNR level and scale the identified time series accordingly to achieve the target SNR.

Since the desired distribution for TCD residuals is non-Gaussian, we used the algorithm presented by Nichols et al. to generate non-white, non-Gaussian time series in step *iii* above, [16]. This algorithm is summarized below:

1. Suppose that the desired number of samples is *n*, and the AR model and generalized normal distribution are known (see steps *i* and *ii*).
2. Generate a realization of the desired generalized normal distribution denoted by *x*_*des-pdf*_ with *n* samples. This realization will likely be white as samples are *Identically and Independently Distributed* (IID).
3. Generate a realization of a normal distribution with *n* samples and filter it using the selected AR model. Denote the filtered normal signal as *x*_*des-AR*_. The filtered signal will also have normal distribution [17]. But, the temporal dynamics would be according to the spectral density of the chosen AR model.
4. Shuffle the samples of *x*_*des-pdf*_ to match the rank of *x*_*des-AR*_, i.e. the smallest value of *x*_*des-pdf*_ is given the same position as the smallest value of *x*_*des-AR*_, the next smallest value of *x*_*des-pdf*_ is given the same position as the second smallest value of *x*_*des-AR*_, etc.

## 3. Results

### 3.1. Power

Figure 1 demonstrates three typical TCD measurements: a scan with large SNR and two scans with relatively smaller SNRs. In the representative scan for high SNR, the power of the residuals was small, Figure 1(A-1), the accepted pulses were consistent and had small interpulse variability, Figure 1(A-2) that gave rise to the large SNR of 21.0 dB. As the interpulse variability increased, the amplitude of residuals increased and SNR decreased dramatically as shown in the last two examples of Figure 1.

Figure 2 shows a typical CBFV measurement and the identified residuals (SNR=3.6 dB). The amplitude structure of the residuals shows that they were well approximated by a generalized normal distribution, Figure 2(B). Besides the location and scale parameters, the generalized normal distribution has the shape parameter *β*. The normal distribution is a special case of the generalized normal distribution with *β* = 2, the tails become heavier than normal for *β* < 2 or lighter than normal for *β >* 2. The shape parameter for this scan was *β* = 1.2. The normal distribution fit is also superimposed for comparison purposes. Since the tails are heavier than a normal distribution, there is frequent outlier looking values in the residuals as evident in Figure 2(A). The power spectral densities of the identified residuals are calculated using Welch’s method in Figure 2(C). They show that the power was larger in the low-frequency region.

**Figure 2:**
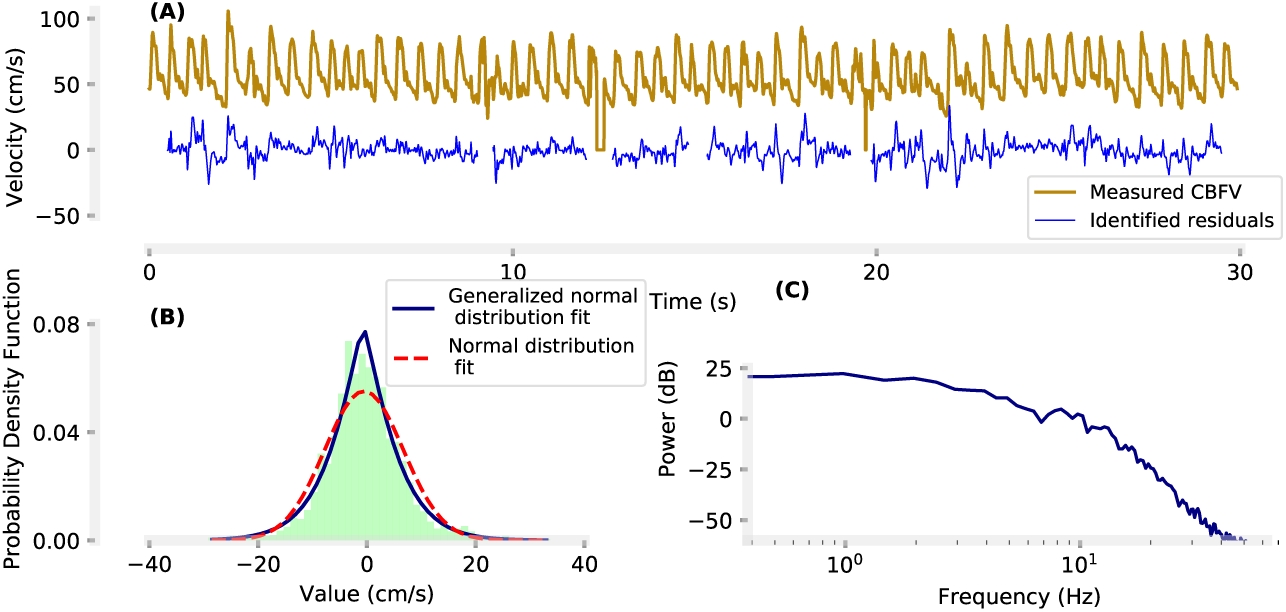
Typical measured CBFV and the identified residuals (A); the amplitude structure of the identified residuals along with the normal and generalized normal parametric fits (B); power spectral density of the identified residuals along with that of the measured CBFV.

Figure 3 shows the distribution of the identified SNRs from the 2567 scans; the mean and standard deviation were 10.5 dB and 5.2 dB, respectively.

**Figure 3:**
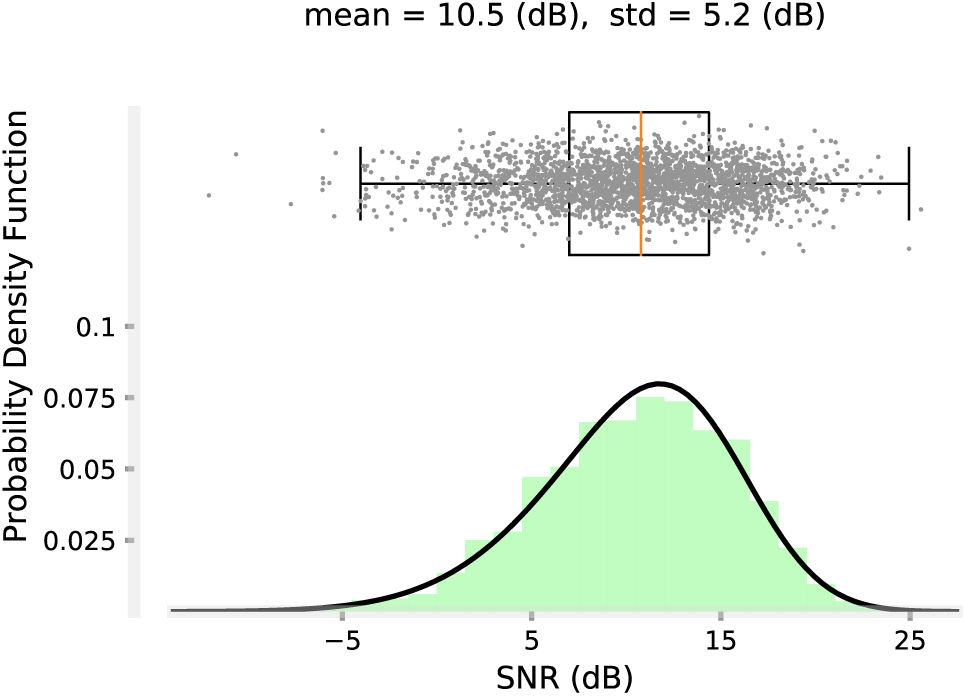
Probability density function of the identified SNRs. The SNR distribution was parameterized by a skewed normal distribution.

### 3.2. Amplitude Structure

We fit theoretical distributions to the empirical distributions identified from the residuals. The generalized normal distribution had the highest quality of fit, quantified in terms of the correlation between the empirical and theoretical distributions, Figure 4. The correlation was larger than 0.9 in more than 96% of the TCD scans. Figure 5 depicts the distribution of the identified *β* parameter. The probability of *β* < 2 was 0.93. Thus, for the majority of the TCD scans, the residuals had a tail heavier than the normal distribution.

**Figure 4:**
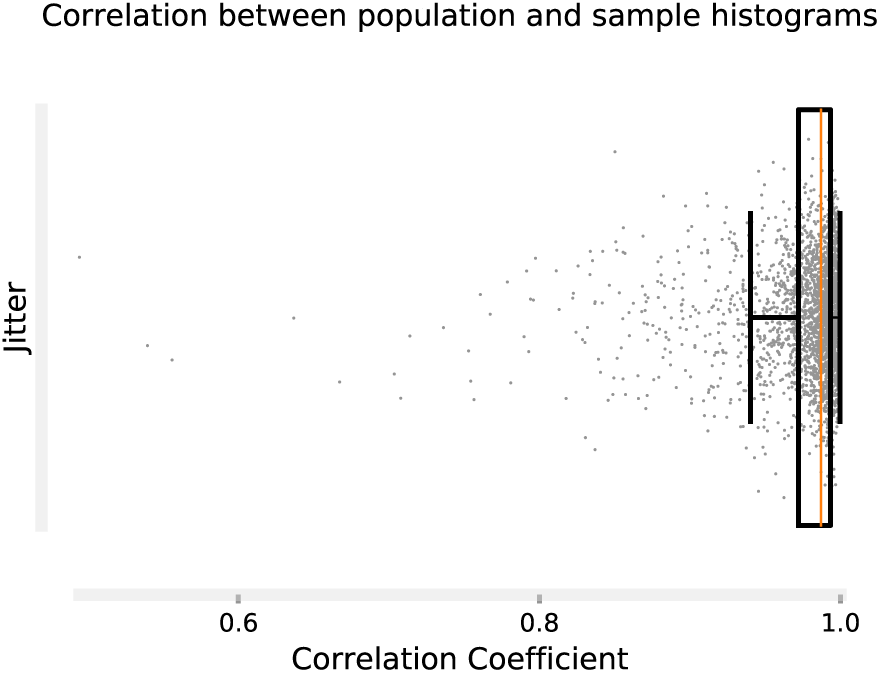
The correlation coefficient between sample empirical PDF and the generalized normal distribution fit. In more than 96% of the TCD scans, the correlation was larger than 0.9 confirming the choice of the generalized normal distribution as an accurate distribution model for TCD residuals.

**Figure 5:**
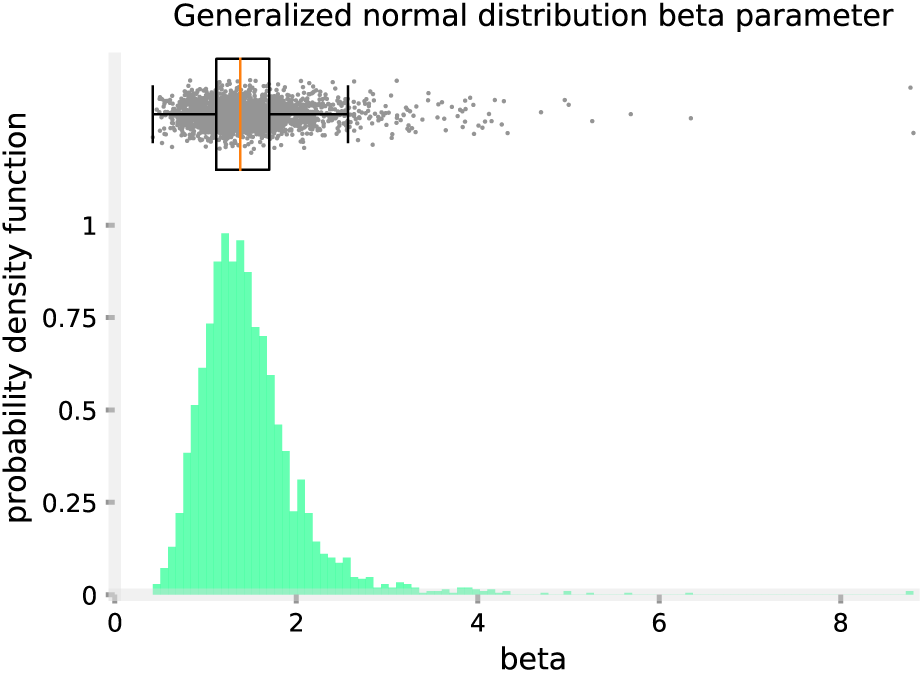
The identified shape parameter *β* of the generalized normal distribution fit to the TCD residuals had a skewed normal distribution. For the majority of the scans (93%), *β* was smaller than 2, demonstrating that the generalized normal distribution fit had tails heavier than normal distribution.

### 3.3. Temporal Structure

50 tap AR models fit the TCD residuals well as confirmed with the analysis of the AR model residuals (not to be confused with the TCD residuals). Figure 6 shows the *Auto-Correlation Coefficient* (ACC) of AR model residuals for a typical TCD scan. Similarly, the ACC of all AR model residuals was one at zero lag and were zero elsewhere demonstrating that they were white, thus confirming the validity of the AR model choice.

**Figure 6:**
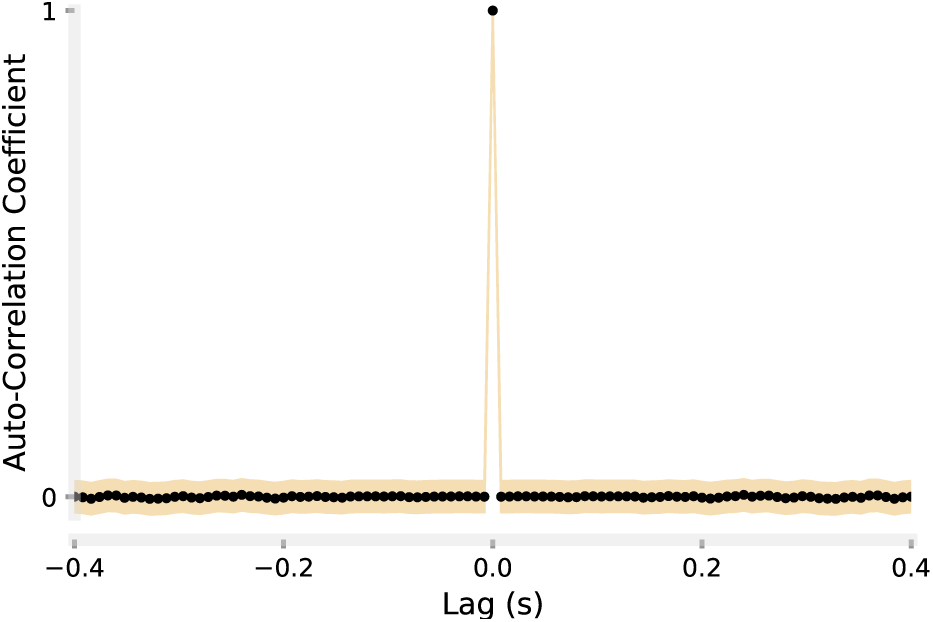
*Auto-Correlation Coefficient* (ACC) of the residuals of a typical AR model. ACC had a spike at lag zero and was zero elsewhere demonstrating that the residuals where white, confirming the validity of the AR model.

Figure 7(A) illustrates the identified 2567 AR model fits to the TCD residuals. For visualization purposes, the magnitude and phase portions of the frequency response function of the identified AR models are shown in Figure 7(B, C) and demonstrate that these models were low-pass in nature with an average break frequency of around 0.5 Hz.

**Figure 7:**
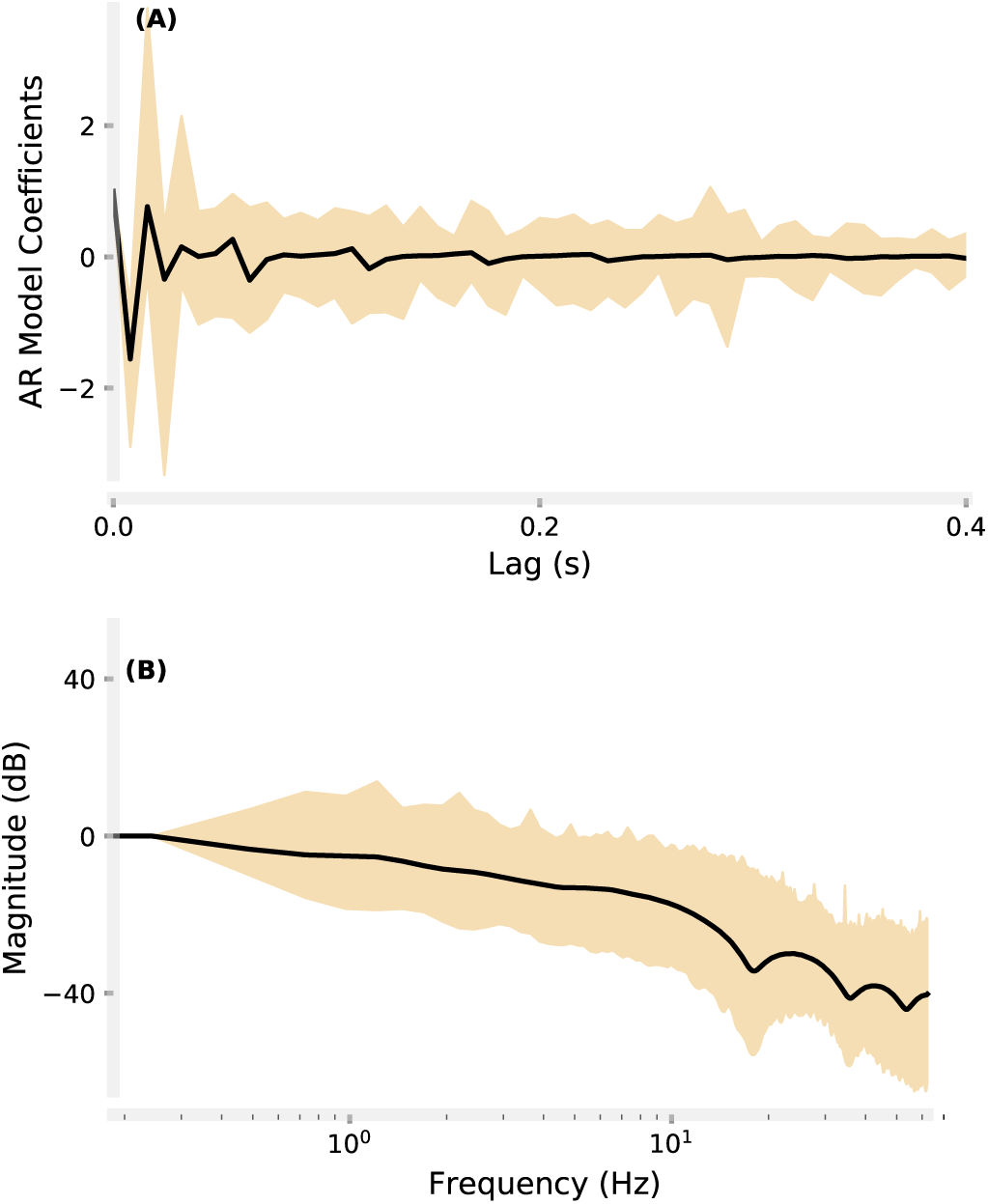
Identified 2567 Auto-Regressive (AR) models: (A) coefficients of the AR models; (B) magnitude portion of the frequency response function representation of the AR models; (C) phase portion of the frequency response function representation of the AR models.

### 3.4. Synthetic TCD Generation

To generate synthetic TCD data, we scaled a pulse archetype (e.g. normal, stenotic, dampened, blunted, etc. [18]) spatially and temporally such that its morphological points lie within the typically observed ranges. We used the following ranges: 3 cm/s < diastolic velocity < 106 cm/s, 7 cm/s < velocity range < 182 cm/s, 42 bpm < heart rate < 151 bpm. Next, we generated the clean TCD signal by tiling copies of the scaled pulse. We simulated synthetic TCD residuals, and scaled it to match the desired SNR level and summed with the clean signal. Figure 8 depicts four examples of synthetic TCD signal with different SNR levels together with their residuals. The experimental and synthetic TCD signals look qualitatively similar demonstrating the potential utility of this tool.

**Figure 8:**
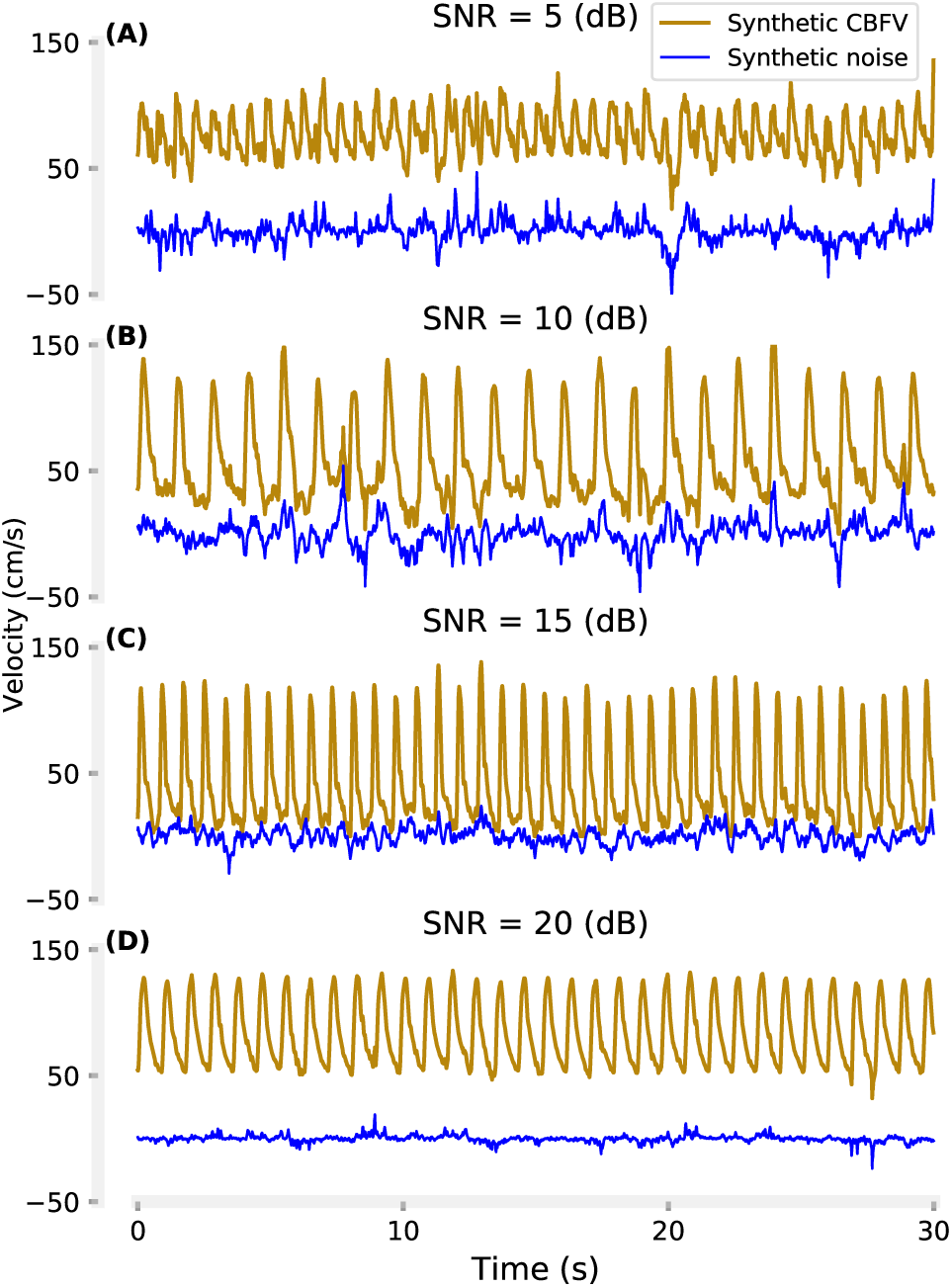
Synthetic TCD signals: (A) SNR=5 dB, (B) SNR=10 dB, (C) SNR=15 dB, (D) SNR=20 dB.

## 4. Discussion

### 4.1. Summary

In this paper, we presented an algorithm to extract TCD residuals. We parameterized the amplitude and temporal structures of the TCD residuals from a rich dataset and identified that the amplitude structure was best characterized by a generalized normal distribution whose tails were heavier than normal distribution for 93% of the scans. For temporal structure, auto-regression models fit the residuals well. They showed that the dynamics were low-pass in nature. Finally, we presented an algorithm to create synthetic TCD residuals that when summed with stereotypical TCD waveforms, resulted in synthetic signals that looked qualitatively similar to experimentally recorded TCD signals.

### 4.2. Signal to Noise Ratio

Analysis of the TCD residuals revealed that our data had a large spectrum of quality, quantified as the SNR. The larger the SNR, the lower the CBFV variability, and the higher the signal quality. It is important to clarify what we mean by “noise” and “signal” in our calculation of SNR. Here, noise simply refers to the TCD residuals, inter-pulse variabilities that degrade the estimate of biomarkers calculated from the average pulse waveform. “Signal” refers to the clean TCD. Since there is no closed-form expression for TCD pulse waveform, we opted to use an empirical model to calculate the clean TCD. We assumed that the vascular activity was in steady-state and we were only interested in the parameters estimated from the average pulse waveform in the rest condition. Thus, any deviation from the average pulse waveform was considered as residual that can effectively alter the confidence in the estimate of the biomarker of interest. Obviously, such an assumption must be revisited according to the experimental protocol, for example, when we present subjects with vasoactive stimuli to test the cerebrovascular reactivity [3, 19]. As such, we need to revisit our definition of the pulse model and accordingly what deviation from the model means, for example by incorporating pulse level variability in heart rate and/or systole/diastole in the empirical model of the pulse waveform.

The identified range for SNR was quite large for our dataset: 90% range was [1.7, 18.2] dB. There are external factors contributing to the signal quality like the ambient noise or movement of the TCD probe/patient. There are also anatomical/(patho)physiological factors such as the thickness of the temporal bone through which we insonate the artery, dampened signal amplitude because of the vessel occlusion, pathologic interpulse variability in TCD pulses (e.g. heart arrhythmia), sharp turns in the artery, etc [20, 4, 21]. Consequently, due to the high number of subjects, both healthy and pathologic, we expect to see such large variability in these factors and subsequently in the SNR.

### 4.3. Algorithm

Time series including sampled CBFV are typically comprised of deterministic and stochastic components. Our algorithm first identified the deterministic component and took the TCD residuals as the stochastic component. Since we acquired TCD during the rest condition, we assumed quasi-steady-state condition and averaged the pulse waveforms to identify the deterministic component as the *Average Pulse Waveform* (APW). Therefore, the stochastic component, i.e. the difference between the acquired data and the deterministic component, represented the deviation from the APW. As explained in 4.2, more complex models must be considered in the presence of vasoactive stimuli or when the scan is long violating the steady-state condition.

In many studies of biomedical signals and systems, “noise” is typically modeled as a white, normal signal. We identified that the TCD residuals were neither white nor normal. Thus, we characterized both amplitude and temporal structures of the stochastic component of TCD. We used probability density function for amplitude structure and identified that the best distribution was generalized normal. We used autoregressive models for temporal structure and identified that they were low-pass in nature.

For simulation of synthetic residuals, the goal is to generate realizations of random sequences conforming to the identified models for distribution and spectral density. Simulation of white, non-normal sequences is straightforward by drawing random variates from the available generators. Since the draws are IID, the resulting sequences will likely be white. Similarly, simulation of non-white normal sequences is also straightforward. One can achieve the target dynamics by passing a white normal sequence through an appropriate linear filter as they preserve the normality [22]. Simulation of non-normal, non-white sequences, is, however, not as straightforward as filtering does not preserve non-normal distribution. This is mainly because filtering is effectively the weighted sum of lagged values of the sequence. Thus, if the sequence has a non-normal distribution, the filter output, which is the sum of non-normal sequences, tends to be close to a normal distribution that can be explained by the central limit theorem [23].

The standard method for generating non-white, non-normal sequences is to first create the desired spectral density by filtering a normal sequence and then to modify its distribution by passing it through a static nonlinearity. The inverse of the desired distribution is used to design the static nonlinearity [24]. Since the closed-form of the inverse of this transformation is typically not available, some numerical methods have been developed to achieve this. One approach is to approximate this numerically by solving many integrals, each of which depends on the calculation of the inverse. This approach is highly complex and computationally demanding [25]. Other data-driven approaches have been proved to be more effective. First, a realization with the desired distribution is generated. Shuffling the samples changes the spectral density of the sequence but does not change the distribution. Stochastic shuffling, i.e. blind shuffling technique, was first proposed by Hunter and Kearney [26]. Later, Nichols et al. improved this approach by using an auxiliary sequence with normal distribution and the desired spectral density that is easy to generate [16]. They used the auxiliary sequence to guide the shuffling process. We adopted this approach for simulation of the TCD residuals as it was fast, simple to implement and showed similar performance to the more complex standard approach.

It is important to make a distinction between *synthetic TCD residuals* and *synthetic TCD signal*. We assumed an additive model and took the synthetic TCD signal as the sum of the synthetic TCD residuals and clean TCD signal. We made a simplifying assumption in the method of generating clean TCD signal that the vascular system was in perfect steady-state, and the resulting CBFV was periodic. This assumption enabled us to simply tile copies of a scaled pulse archetype to build the clean TCD signal. We did not make an attempt to simulate the physiological or pathophysiological variability in the synthetically generated TCD pulses. Note that the algorithm to generate *synthetic TCD residuals* is general in the sense that it simulates stochastic time series conforming to arbitrary models for amplitude and temporal structures and does not hinge on the simplifying assumption of the pulse waveform model.

### 4.4. Clinical Implication

The proposed algorithm provides a single, easy to use, meaningful metric, i.e. SNR, to assess signal quality following a TCD scan that can be implemented as part of a post hoc analysis routine. The SNR is a normalized metric describing the relative power of the residuals with respect to the signal (modeled TCD signal). To put the identified SNR values from our experimental results in perspective, a value larger than 0 dB indicates that the signal power is larger than noise power, 10 dB indicates that the signal power is 10 times larger than noise power, and 20 dB indicates that the signal power is 100 times larger than noise power and so on. Based on the acceptable tolerance level, this metric can facilitate the decision making for clinicians and TCD operators on whether to keep the acquired data following a scan for diagnostic purposes or discard and search for a higher quality signal. SNR can also be used as an ultimate metric to objectively compare TCD probes, signal processing algorithms, electronics, etc in practical tests.

The algorithm for the generation of synthetic TCD residuals is useful to study the uncertainty associated with the estimate of biomarkers. Monte-Carlo simulations can be designed to characterize the error in a range of expected SNR levels and quantify the confidence level. This algorithm is also useful in creating a realistic TCD emulation platform that can be used to train neurosonographers.

### 4.5. Areas of future work

To the best of our knowledge, for the first time, we parameterized the temporal structure of the TCD residuals. As a first step, we used simple linear AR models to capture the dynamics of the TCD residuals. The AR models can only capture stationary stochastic components. In the future, it would be within our interest to test more complex models. Time-varying or nonlinear AR/ARMA, or multiplicative models can be more effective to capture a range of possible non-stationarities and nonlinearities such as modulation of the residual power with CBFV pulses.

The variabilities that constitute the TCD residuals can have intrinsic origins, e.g. heart rate variability, auto-regulatory mechanisms, or extrinsic origins such as movement artifacts, electronics, etc. As a first attempt to characterize the TCD residuals we studied these lumped together. If there is interest to study these components individually, more complex models of the pulse waveform are needed to extract the variabilities with physiological origins.

The method presenting in this work is general and would be a powerful tool for other biomedical systems to characterize additive residual variance. The key assumption is that the deterministic component of the system is identifiable to subtract from the measured noisy signal to give the stochastic component. Characterization of the deterministic component generating quasi-periodic signals including the TCD waveform studied here is straightforward using data-driven tools (e.g. ensemble averaging used here). Other examples of quasi-periodic signals include measurement of arterial blood pressure, ECG, respiration signals, physiological and pathological tremors. For other systems, the deterministic component can be identified in a separate experiment by manipulating the input signals and perturbing the system such as models of muscle spindles, functional electrical stimulation, etc. [27, 28]. In other systems, the input signals can be set to be zero to remove the contribution of the deterministic component altogether, for example, eye movement or joint torque during rest [29, 30].

## 6. Acknowledgment

The authors thank Dr. James K. Fleming, Ms. Brenda Knowles, Mr. Leo Martinez, Mr. Ben McClellan, Ms. Jennifer Nichols, Ms. Jennifer Patterson, and Dr. Ruchir Shah for their help during data collection and subject recruitment.

